# Immunogenicity of low dose prime-boost vaccination of mRNA vaccine CV07050101 in non-human primates

**DOI:** 10.1101/2021.07.07.451505

**Authors:** Neeltje van Doremalen, Robert J. Fischer, Jonathan E. Schulz, Myndi G. Holbrook, Brian J. Smith, Jamie Lovaglio, Benjamin Petsch, Vincent J. Munster

**Affiliations:** Laboratory of Virology, National Institute of Allergy and Infectious Diseases, National Institutes of Health, Hamilton, MT, USA; Rocky Mountain Veterinary Branch, National Institute of Allergy and Infectious Diseases, National Institutes of Health, Hamilton, MT, USA; CureVac AG, Tuebingen, Germany

## Abstract

Many different vaccine candidates against severe acute respiratory syndrome coronavirus-2 (SARS-CoV-2), the etiological agent of COVID-19, are currently approved and under development. Vaccine platforms vary from mRNA vaccines to viral-vectored vaccines, and several candidates have been shown to produce humoral and cellular responses in small animal models, non-human primates and human volunteers. In this study, six non-human primates received a prime-boost intramuscular vaccination with 4 µg of mRNA vaccine candidate CV07050101, which encodes a pre-fusion stabilized spike (S) protein of SARS-CoV-2. Boost vaccination was performed 28 days post prime vaccination. As a control, six animals were similarly injected with PBS. Humoral and cellular immune responses were investigated at time of vaccination, and two weeks afterwards. No antibodies could be detected two and four weeks after prime vaccination. Two weeks after boost vaccination, binding but no neutralizing antibodies were detected in 4 out of 6 non-human primates. SARS-CoV-2 S protein specific T cell responses were detected in these 4 animals. In conclusion, prime-boost vaccination with 4 µg of vaccine candidate CV07050101 resulted in limited immune responses in 4 out of 6 non-human primates.

## Introduction

Severe acute respiratory syndrome coronavirus 2 (SARS-CoV-2) is the etiological agent responsible for COVID-19. SARS-CoV-2 has spread worldwide and over 185 million cases have been detected as of July 2021. The pandemic has resulted in an unprecedented research effort towards the development of a SARS-CoV-2 vaccine and several vaccines against SARS-CoV-2 have now been approved. Interestingly, whilst traditional approaches such as subunit protein vaccines^1^ and inactivated virus vaccines^2^ are still pursued, a large number of vaccines are based on novel platforms, such as virus-vectored vaccines^3-5^ and nucleic acid (DNA or RNA) vaccines^6,7^. Promising results have been published for these platforms, both preclinical^8-13^ and clinical^3-7^, showing the induction of a humoral and cellular response.

Preclinical assessment of SARS-CoV-2 vaccines in non-human primate models is advantageous due to the close relatedness of non-human primates to humans, thereby resulting in a higher degree of clinical translation than smaller animal models. Indeed, rhesus macaques have been successfully used to study vaccines^14^. Inoculation of rhesus macaques with SARS-CoV-2 results in respiratory disease which includes virus replication in upper and lower respiratory tract^15^. Two reports on the immune response of SARS-CoV-2 mRNA vaccine candidates in non-human primates describe the induction of binding and neutralizing antibodies, as well as antigen-specific T cell responses^9,10^.

SARS-CoV-2 messenger RNA (mRNA) vaccines encoding the SARS-CoV-2 spike (S) protein have a good safety and immunogenicity profile, both in non-human primates^9,10^ and in humans^6,7,16^. Here, we investigate the immunogenicity of another SARS-CoV-2 S mRNA vaccine, CV07050101, in non-human primates. CV07050101 is based on mRNA technology, RNActive^®^, developed by CureVac for the accelerated development of human vaccines^17-21^. The efficaciousness of this platform has been demonstrated for a rabies vaccine in mice and humans^18,22^. Moreover, mRNA vaccines have been discussed as particular well suited to combat outbreak pathogens^23^.

## Results

In order to investigate the immunogenicity of mRNA vaccine CV07050101, we vaccinate six rhesus macaques (all male) at 0 and 28 days via intramuscular injection, using 4 µg per dose. As a control, six rhesus macaques were injected with an equal volume of sterile PBS (Figure 1A). No adverse events were observed upon vaccination, and overall hematology and clinical chemistries were unremarkable. No differences between the control and vaccinated groups were noted. No binding antibodies could be detected 14 or 28 days post prime vaccination (Figure 1B). 14 days post boost vaccination, low titers of spike-specific binding antibodies (reciprocal endpoint IgG titers of 400-800) could be detected in 4 out of 6 animals (Figure 1B). Virus-specific neutralizing antibodies were not detected in animals at any time post boost vaccination (Figure 1C). No SARS-CoV-2 spike-specific T cell responses were detected 14 days post prime vaccination but were detected in the same 4 out of 6 animals at 14 days post boost vaccination (Figure 1D). The detection of specific T cell responses correlated with the detection of spike-specific binding antibodies.

**Figure.**
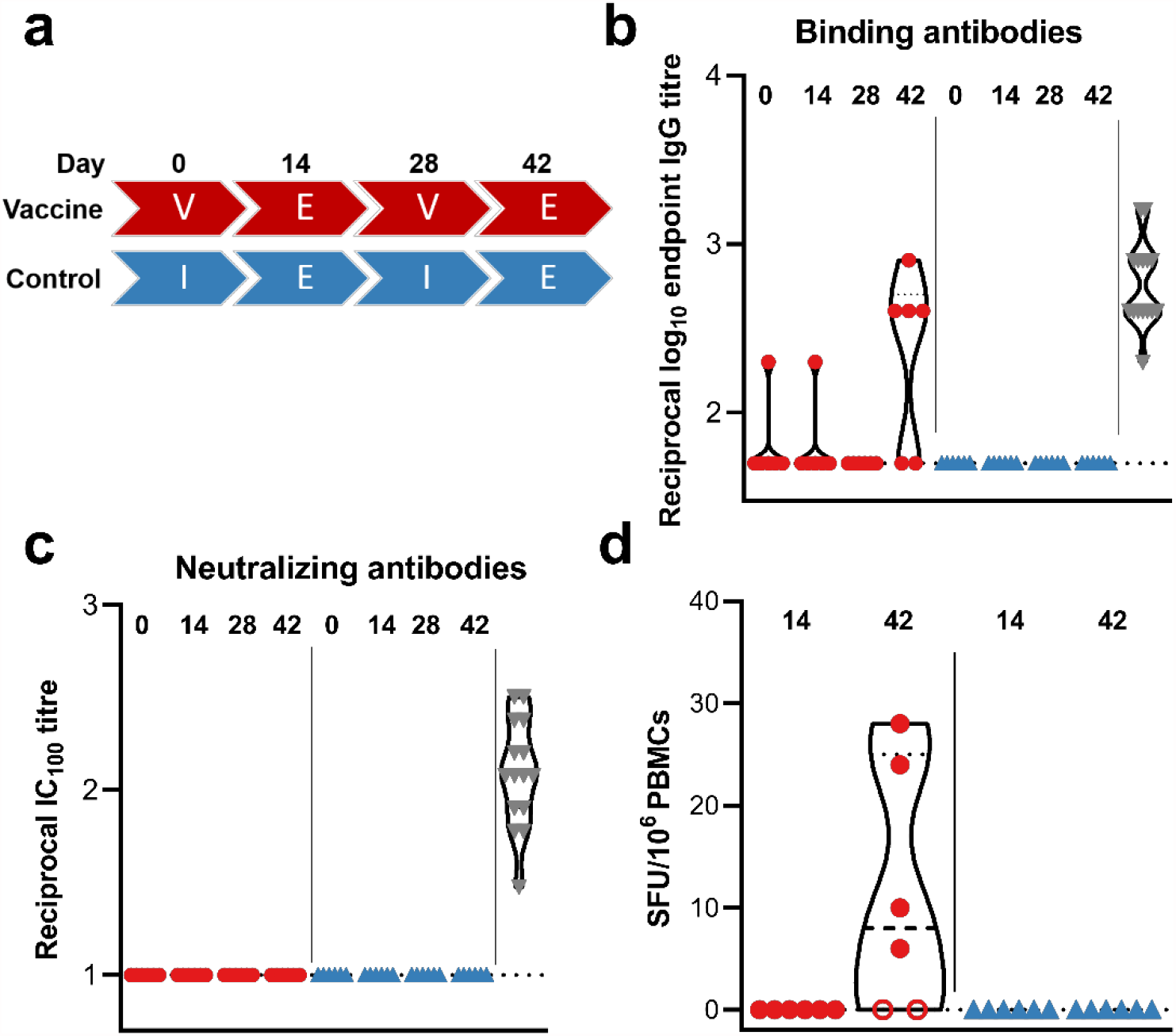
Humoral and cellular response after vaccination with CV07050101. a) Study schedule. Two groups (N=6) were vaccinated (V) or administered PBS (I) twice, four weeks apart. Fourteen days post each vaccination, exams (E) were performed. The presence of SARS-CoV-2 spike-specific binding (b) and neutralizing (c) antibodies in serum obtained from rhesus macaques at time of vaccination and 14 days afterwards where measured using ELISA and infectious virus neutralization assays. d) SARS-CoV-2 S-specific T cell responses were measured via ELIspot. Closed red circles = animal positive in ELISA assay; open red circle = animal negative in ELISA assay; blue triangles = control animals; grey triangles = convalescent human sera; dotted line = lower limit of detection.

## Discussion

Here, we show that prime-boost vaccination of rhesus macaques with 4 µg of CV07050101 results in the induction of binding antibodies in some, but not all vaccinated animals. This contrasts with other studies with mRNA vaccines, in which a prime-boost vaccination elicits a robust humoral and cellular response in all animals. Using a prime-boost regimen of 10 µg of mRNA-1273, which encodes a prefusion-stabilized S protein utilizing modified mRNA, S-specific binding antibodies were detected in all animals, whereas neutralizing antibodies were detected in 7 out of 8 animals^9^. Likewise, a prime-boost vaccination using 30 µg of vaccine candidate BNT162b2, which also encodes a prefusion-stabilized S protein, elicits binding and neutralizing antibodies in 6 out of 6 animals. Moreover, binding and neutralizing antibodies were detected in all animals 21 days post prime only vaccination with 30 µg of BNT162b2^10^. A recently released preprint investigating CV07050101 showed that vaccination of rhesus macaques with 8 µg of mRNA elicits binding and neutralizing antibodies, whereas a dose of 0.5 µg of mRNA did not^23^.

One important difference between these studies is the amount of mRNA used to vaccinate animals. We used 4 µg of mRNA per vaccination, whereas the other doses which elicited an immune response used between 8 µg and 100 µg per vaccination^9,10,24^. Using a dose of 10 µg mRNA-1273 vaccine resulted in no detectable neutralizing antibodies in 1 out of 8 animals on the day of challenge, but 100 µg of mRNA-1273 resulted in neutralizing antibodies in all animals^9^, suggesting a dose-dependent response to the vaccine. Likewise, whereas 8 µg of CV07050101 induced an immune response, 0.5 µg did not^23^.

Compared to the limited immunogenicity in non-human primates we observed here, robust SARS-CoV-2 neutralizing titers were observed in Balb/c mice immunized with the CV07050101 vaccine after prime-boost regimen. Challenge studies in hamsters, which were performed at a later stage, utilized a 10 µg prime-boost regimen of CV07050101 vaccine and a challenge dose of 10^2^ TCID_50_ SARS-CoV-2 and provided protection of the lower respiratory tract^24^.

As the elicited immune response was low or absent in the vaccinated rhesus macaques, we decided not to challenge the animals. In rhesus macaques, neutralizing antibodies are a correlate of protection^25^. The presence of neutralizing antibodies in humans correlates with immunity against SARS-CoV-2^26^. Since we did not detect neutralizing antibodies, we hypothesize that these animals would not have been protected.

Since CV07050101 is now assessed in clinical trial studies, we compared the available immunogenicity results. The 12 µg high dose vaccine prime-boost regime was able to induce neutralizing antibody titers comparable to non-hospitalized individuals, whereas the 2-8 µg doses induced neutralizing titers that were lower in clinical trial participants. However, virus neutralizing antibodies could be detected in 66% of human volunteers given 4 µg of CV07050101^16^, in contrast to no neutralizing antibodies in serum from NHPs vaccinated with the same dose. It has been hypothesized that a difference in the lubricant used in syringes can decrease integrity of the mRNA vaccine, which may explain the low immunogenicity detected in this NHP study (B.P. personal communication).

In conclusion, we show that prime-boost vaccination of rhesus macaques with 4 µg of CV07050101 does not elicit a uniform nor robust immune response. However, vaccination using 8 µg of the same vaccine was protective in a NHP challenge study demonstrating protection against SARS-CoV-2 infection by CV07050101 vaccination^23^.

## Methods

### Ethics statement

Animal study approval was provided by the Institutional Animal Care and Use Committee (IACUC) at Rocky Mountain Laboratories. Animal experiments were conducted in an AAALAC-approved facility, following the basic principles and guidelines in The Guide for the Care and Use of Laboratory Animals, the Animal Welfare Act, United States Department of Agriculture and the United States Public Health Service Policy on Humane Care and Use of Laboratory Animals. Rhesus macaques were housed in individual primate cages allowing social interactions, in a climate-controlled room with a fixed light/dark cycle (12-hours/12-hours). Animals were monitored a minimum of twice daily and commercial monkey chow, treats, vegetables, and fruit were provided. Water was available ad libitum. A variety of human interaction, commercial toys, videos, and music was used as environmental enrichment.

### Vaccine mRNA and lipid nanoparticle production

CV07050101 is an lipid nanoparticle-formulated RNActive® SARS-CoV-2 vaccine composed of the active pharmaceutical ingredient, an mRNA that encodes a pre-fusion conformation stabilized version of the full-length spike (S) protein of SARS-CoV-2 virus (GenBank YP_009724390.1) including the K986P and V987P prefusion stabilizing mutations, and four lipid components: cholesterol, 1,2-distearoyl-sn-glycero-3-phosphocholine (DSPC), PEGylated lipid and a cationic lipid^24^.

### Study design

12 male rhesus macaques between 3-5 years old were screened for SARS-CoV-2 status by ELISA, and when found to be negative for prior exposure were sorted by body weight, and then divided into two groups of six animals, resulting in near equal contribution of body weights. Group 1 (vaccine) was vaccinated with 4 µg of mRNA vaccine CV07050101 in sterile PBS at 0 and 28 days, group 2 (control) was vaccinated with sterile PBS at 0 and 28 days via intramuscular injection, using Monoject 1 mL Tuberculin syringes (Covidien, 25G x 5/8”). Blood samples were obtained before vaccination and 14 days after each vaccination. Hematology analysis was completed on a ProCyte DX (IDEXX Laboratories, Westbrook, ME, USA) and the following parameters were evaluated: red blood cells (RBC), hemoglobin (Hb), hematocrit (HCT), mean corpuscular volume (MCV), mean corpuscular hemoglobin (MCH), mean corpuscular hemoglobin concentration (MCHC), red cell distribution weight (RDW), platelets, mean platelet volume (MPV), white blood cells (WBC), neutrophil count (abs and %), lymphocyte count (abs and %), monocyte count (abs and %), eosinophil count (abs and %), and basophil count (abs and %). Serum chemistries were completed on a VetScan VS2 Chemistry Analyzer (Abaxis, Union City, CA) and the following parameters were evaluated: glucose, blood urea nitrogen (BUN), creatinine, calcium, albumin, total protein, alanine aminotransferase (ALT), aspartate aminotransferase (AST), alkaline phosphatase (ALP), total bilirubin, globulin, sodium, potassium, chloride, and total carbon.

### Enzyme-linked immunosorbent assay

A plasmid encoding the prefusion stabilized SARS-CoV-2 spike protein with a T4 fibritin trimerization motif was obtained from the Vaccine Research Centre, Bethesda, USA and expressed in-house. Maxisorp plates (Nunc) were coated overnight at 4 °C with 100 ng/well spike protein in PBS. Plates were blocked with 100 µl of casein in PBS (Thermo Fisher) for 1hr at RT. Serum serially diluted 2x in casein in PBS was incubated at RT for 1hr. Antibodies were detected using affinity-purified polyclonal antibody peroxidase-labeled goat-anti-monkey IgG (Seracare, 074-11-021) in casein and TMB 2-component peroxidase substrate (Seracare, 5120-0047), developed for 5-10 min, and reaction was stopped using stop solution (Seracare, 5150-0021) and read at 450 nm. All wells were washed 3x with PBST 0.1% tween in between steps. Threshold for positivity was set at 3x OD value of negative control (serum obtained from non-human primates prior to start of the experiment) or 0.2, whichever one was higher.

### ELISpot

PBMCs were isolated from ethylene diamine tetraaceticacid (EDTA) whole blood using LeucosepTM tubes (Greiner Bio-one International GmbH) and Histopaque®-1077 density gradient cell separation medium (Sigma-Aldrich) according to the manufacturers’ instructions. IFN-γ ELISpot assay of PBMCs was performed using the ImmunoSpot® Human IFN-γ Single-Color Enzymatic ELISpot Assay Kit according to the manufacturer’s protocol (Cellular Technology Limited). PBMCs were plated at a concentration of 300,000 cells per well and were stimulated with two contiguous peptide pools spanning the length of the SARS-CoV-2 S protein sequence at a concentration of 2 µg/mL per peptide (Mimotopes). Imaging was performed using the CTL ImmunoSpot® Software (Cellular Technology Limited). Spot forming units (SFU) were hand counted and calculated per 10^6^ PBMCs as summed across the peptide pools for each animal.

### SARS-CoV-2 virus neutralization

VeroE6 cells were maintained in Dulbecco’s modified Eagle’s media (DMEM) supplemented with 10% fetal bovine serum (Gibco), 1 mM L-glutamine, 50 U/mL streptomycin and 50 ug/mL penicillin. Sera were heat-inactivated (30 min, 56 °C), two-fold serial dilutions were prepared in DMEM supplemented with 2% fetal bovine serum (Gibco), 1 mM L-glutamine, 50 U/mL streptomycin and 50 ug/mL penicillin and 100 TCID_50_ of SARS-CoV-2 was added. After 1hr incubation at 37 °C and 5% CO_2_, virus:serum mixture was added to VeroE6 cells and incubated at 37 °C and 5% CO_2_. At 6 dpi, cytopathic effect was scored. The virus neutralization titer was expressed as the reciprocal value of the highest dilution of the serum which still inhibited 100% of virus replication. A positive control standardized against the NIBSC serum control 20/130 was used in all VN assays.

## Acknowledgements

We thank Olubukola Abiona, Victoria Avanzato, Kaitlyn Bauer, Chase Baune, Kizzmekia Corbett, Kathleen Cordova, Shane Gallogly, Barney Graham, Brian Hancock, Patrick Hanley, Corey Henderson, Billy Jameson, Michael Jones, Rachel LaCasse, Kay Menk, Jyothi Purushotham, Rocky Rivera, Jeff Severson, Les Shupert, and Marissa Woods for their assistance during this study.

## Author Contributions

N.v.D and V.M. designed the studies, N.v.D, R.J.F., J.S., M.G.H., B.J.S., and J.L. performed the studies, B.P. provided the vaccine, N.v.D. analyzed the results, N.v.D. wrote the manuscript, all co-authors reviewed the manuscript.

## References

1. Keech C, Albert G, Cho I, et al. Phase 1-2 Trial of a SARS-CoV-2 Recombinant Spike Protein Nanoparticle Vaccine. N Engl J Med 2020.

2. Zhang Y, Zeng G, Pan H, et al. Immunogenicity and Safety of a SARS-CoV-2 Inactivated Vaccine in Healthy Adults Aged 18-59 years: Report of the Randomized, Double-blind, and Placebo-controlled Phase 2 Clinical Trial. MedRxiv 2020.

3. Folegatti PM, Ewer KJ, Aley PK, et al. Safety and immunogenicity of the ChAdOx1 nCoV-19 vaccine against SARS-CoV-2: a preliminary report of a phase 1/2, single-blind, randomised controlled trial. Lancet 2020;396:467–78.

4. Logunov DY, Dolzhikova IV, Zubkova OV, et al. Safety and immunogenicity of an rAd26 and rAd5 vector-based heterologous prime-boost COVID-19 vaccine in two formulations: two open, non-randomised phase 1/2 studies from Russia. Lancet 2020.

5. Zhu FC, Li YH, Guan XH, et al. Safety, tolerability, and immunogenicity of a recombinant adenovirus type-5 vectored COVID-19 vaccine: a dose-escalation, open-label, non-randomised, first-in-human trial. Lancet 2020;395:1845–54.

6. Jackson LA, Anderson EJ, Rouphael NG, et al. An mRNA Vaccine against SARS-CoV-2 - Preliminary Report. N Engl J Med 2020.

7. Mulligan MJ, Lyke KE, Kitchin N, et al. Phase 1/2 study of COVID-19 RNA vaccine BNT162b1 in adults. Nature 2020.

8. van Doremalen N, Lambe T, Spencer A, et al. ChAdOx1 nCoV-19 vaccine prevents SARS-CoV-2 pneumonia in rhesus macaques. Nature 2020.

9. Corbett KS, Flynn B, Foulds KE, et al. Evaluation of the mRNA-1273 Vaccine against SARS-CoV-2 in Nonhuman Primates. N Engl J Med 2020.

10. Vogel AB, Kanevsky I, Che Y, et al. A prefusion SARS-CoV-2 spike RNA vaccine is highly immunogenic and prevents lung infection in non-human primates. bioRxiv 2020:2020.09.08.280818.

11. Mercado NB, Zahn R, Wegmann F, et al. Single-shot Ad26 vaccine protects against SARS-CoV-2 in rhesus macaques. Nature 2020.

12. Graham SP, McLean RK, Spencer AJ, et al. Evaluation of the immunogenicity of prime-boost vaccination with the replication-deficient viral vectored COVID-19 vaccine candidate ChAdOx1 nCoV-19. NPJ vaccines 2020;5:69.

13. Corbett KS, Edwards DK, Leist SR, et al. SARS-CoV-2 mRNA vaccine design enabled by prototype pathogen preparedness. Nature 2020.

14. Munoz-Fontela C, Dowling WE, Funnell SGP, et al. Animal models for COVID-19. Nature 2020;586:509–15.

15. Munster VJ, Feldmann F, Williamson BN, et al. Respiratory disease in rhesus macaques inoculated with SARS-CoV-2. Nature 2020;585:268–72.

16. Peter Kremsner PM, Jacobus Bosch, Rolf Fendel, Julian J. Gabor, Andrea Kreidenweiss, Arne Kroidl, Isabel Leroux-Roels, Geert Leroux-Roels, Christoph Schindler, Mirjam Schunk, Thirumalaisamy P. Velavan, Mariola Fotin-Mleczek, Stefan Müller, Gianluca Quintini, Oliver Schönborn-Kellenberger, Dominik Vahrenhorst, Thomas Verstraeten, Lisa Walz, Olaf-Oliver Wolz, Lidia Oostvogels. Phase 1 Assessment of the Safety and Immunogenicity of an mRNA-Lipid Nanoparticle Vaccine Candidate Against SARS-CoV-2 in Human Volunteers. medRXIV 2020.

17. Rauch S, Lutz J, Kowalczyk A, Schlake T, Heidenreich R. RNActive(R) Technology: Generation and Testing of Stable and Immunogenic mRNA Vaccines. Methods Mol Biol 2017;1499:89–107.

18. Lutz J, Lazzaro S, Habbeddine M, et al. Unmodified mRNA in LNPs constitutes a competitive technology for prophylactic vaccines. NPJ vaccines 2017;2:29.

19. Fotin-Mleczek M, Duchardt KM, Lorenz C, et al. Messenger RNA-based vaccines with dual activity induce balanced TLR-7 dependent adaptive immune responses and provide antitumor activity. J Immunother 2011;34:1–15.

20. Petsch B, Schnee M, Vogel AB, et al. Protective efficacy of in vitro synthesized, specific mRNA vaccines against influenza A virus infection. Nat Biotechnol 2012;30:1210–6.

21. Schnee M, Vogel AB, Voss D, et al. An mRNA Vaccine Encoding Rabies Virus Glycoprotein Induces Protection against Lethal Infection in Mice and Correlates of Protection in Adult and Newborn Pigs. PLoS Negl Trop Dis 2016;10:e0004746.

22. Armbruster N, Jasny E, Petsch B. Advances in RNA Vaccines for Preventive Indications: A Case Study of A Vaccine Against Rabies. Vaccines (Basel) 2019;7.

23. Rauch S, Jasny E, Schmidt KE, Petsch B. New Vaccine Technologies to Combat Outbreak Situations. Front Immunol 2018;9:1963.

24. Rauch S, Roth N, Schwendt K, Fotin-Mleczek M, Mueller SO, Petsch B. mRNA-based SARS-CoV-2 vaccine candidate CVnCoV induces high levels of virus-neutralising antibodies and mediates protection in rodents. NPJ vaccines 2021;6:57.

25. McMahan K, Yu J, Mercado NB, et al. Correlates of protection against SARS-CoV-2 in rhesus macaques. Nature 2021;590:630–4.

26. Addetia A, Crawford KHD, Dingens A, et al. Neutralizing Antibodies Correlate with Protection from SARS-CoV-2 in Humans during a Fishery Vessel Outbreak with a High Attack Rate. J Clin Microbiol 2020;58.

